# Bioethanol from Sago Waste Fermented by Baker’s and Tapai Yeast as a Renewable Energy Source

**DOI:** 10.1101/2020.01.03.894691

**Authors:** Muhammad Rijal

## Abstract

Renewable energy is collected through sustainable natural processes. One of the examples of renewable energy is a biofuel. A biofuel is produced from organic materials, such as plants with high sugar content. Sago starch extracted from sago pith contains starch and fiber that can be converted into glucose by hydrolysis. Sago starch and fiber can be processed into bioethanol as the main ingredient of renewable energy source. In this study, bioethanol production from sago waste fermented by baker’s yeast and tapai underwent the following stages: delignification, hydrolysis, fermentation, and distillation. The highest bioethanol content was obtained from the BRT treatment where wet solid sago waste was fermented by baker’s yeast (45.7021%), while the lowest was found in the BTP treatment (0.9504%). Two or more than two peaks were shown by the ART, ATP, BTP, and KTP treatments, whereas only one peak was indicated by the BRT and KRT treatments suggesting that there was only one compound that can be identified as ethanol.

## Introduction

Sago is one of the staples of the Indonesian diet. People in Maluku consume sago on a daily basis. The production of sago into starch results in sago waste in the form of sago fiber and liquid waste which are beneficial for Maluku people in particular, and Indonesian people in general. Solid and liquid sago waste can be processed into renewable and environmentally-friendly energy sources. The sago starch extraction industry commonly produces three types of waste; they are fibrous sago pith residue or sago waste (hampas), sago bark waste, and sago wastewater. Sago bark waste and sago pith waste weigh around 26% and 14% of the total weight of a sago block (Singhal et al., 2008; Idral, Salim, and Mardiah, 2012; Fitriana, 2009). Sago waste is reported to contain important components such as starch and cellulose. According to Fitriana (2009), sago waste is made up of 65.7% starch and 34.3% crude fiber, crude protein, fat, and ash which can be processed into a renewable energy source (Fitriana, 2009).

Renewable energy sources can recover quickly and can be sustained indefinitely. If managed and utilized properly, these energy sources can replace fossil energy in the future. A biofuel is a renewable energy source that is produced from organic materials, such as plants or organic waste that has high sugar content and can be processed into bioethanol (Hambali et al., 2007).

Bioethanol is abbreviated as Et-OH. The molecule formula of ethanol is C_2_H_5_OH, while its empirical formula is C_2_H_6_O and its structural formula is written as CH_3_-CH_2_-OH. Bioethanol is a methyl (CH_3_-) group that is bound to a methylene group (-CH_2_-) and a hydroxyl group (-OH) (Richana, 2011; Riadi, 2013). Research on bioethanol as a renewable energy source and environmentally friendly substitute for fossil fuels has been often conducted, such as bioethanol produced from *kepok* banana peel waste (Setiawati, 2013), bioethanol produced from corn stalk waste (Muniroh 2011), and bioethanol produced from cassava skin waste. (Hikmayati and Yanie, 2008). These three types of waste contain a lot of carbohydrates in the form of starch and lignin and so does sago waste. Therefore, sago waste can be potentially used as a raw material in bioethanol production. Bioethanol production from organic materials that contain high levels of carbohydrate is supported by enzymatic and microorganism activities. The results of a study conducted by Nugraha (2012) showed that on a laboratory scale, the use of 100 gr alpha-amylase, 100 g glucoamylase, and 100 gr yeast produced the highest ethanol content (1.110%) per 1 kg sago starch.

The use of sago starch as a raw material in bioethanol production is considered inefficient because sago starch is the staple of some Indonesian people’s diet. Thus, instead of using sago starch, the people can take an advantage of the sago waste. Bioethanol production using enzymes is also costly; therefore, acid and yeast compounds that are relatively affordable can be used as the alternatives. A study indicated that the composition of sago pith waste and tapai yeast a ratio of 9%: 7% can result in 2.1% ethanol content (Gustari et al., 2012). Sago waste can be found in various forms such as in solid and liquid forms. Solid sago waste consists of wet and dry solid waste.

This study aimed to determine the content and the profile of bioethanol produced from sago pith waste fermented by yeast. Types of sago waste that were used as the source of lignocellulose included wet solid, dry solid, and liquid sago waste. Baker’s yeast and tapai yeast were utilized to ferment the waste and convert it into glucose through an enzymatic process. Accordingly, three research questions were formulated: (1) What are bioethanol profiles produced from sago waste fermented by baker’s yeast and tapai yeast?; (2) What is bioethanol content produced from sago waste fermented by baker’s yeast and tapai yeast?; (3) Do types of sago waste and types of yeast affect bioethanol content?

The current research report contains the following arguments: (1) the profile of bioethanol produced from distilled fermented sago waste varied depending on types of sago waste and types of yeast used in the process (using GCMS to observe bioethanol graphs); (2) sago waste processed with different treatments resulted in different bioethanol content (using a refractometer to test bioethanol levels); (3) Types of sago and types of yeast have an effect on bioethanol content (observing the difference in bioethanol production levels from each treatment).

## Literature Review

### Renewable Energy

The current energy crisis in Indonesia is mainly caused by the depletion of petroleum reserves. Sooner or later, the world’s oil reserves will be exhausted due to their limited and non-renewable supplies. These circumstances have encouraged many countries in the world to increase their efforts to use biofuels as alternative fuels. Biofuels are fuels or energy sources that come from organic materials. Biofuels can be renewed. Therefore, they are identified as renewable energy sources (Trisanti, 2009).

The oil depletion issue is an urgent call to explore alternative energy sources that can be renewed. One of the examples of alternative energy sources is bioethanol. The use of ethanol as bioethanol has several advantages, including reducing the dependency on petroleum, creating jobs in the correspondent area, and reducing air pollution because it minimizes the formation of CO2 gas. The ever-increasing price of fuel oil and the increasingly limited world oil reserves have encouraged efforts to obtain alternative fuels (Zaldivar *et al*., 2001; Schubert, 2006).

### Bioethanol Production

Bioethanol is an alternative energy that is widely produced in the world today. The use of lignocellulosic raw materials in bioethanol production encourages the development of renewable energy businesses and the reduction of fuel oil production costs (Knauf and Moniruzzaman, 2004; Ragauskas *et al*., 2006; Schubert, 2006). The use of these raw materials minimizes fears about competition in plant-based food manufacture. Biomass used as a raw material to produce energy also plays a crucial role in reducing greenhouse gas emissions because CO2 released from the degradation of natural biomass will be available as carbon in energy (Schubert, 2006).

The process of creating bioethanol from sago waste should go through the following stages: delignification, hydrolysis, and fermentation. Delignification is the simplification of lignin using acidic compounds. Delignification is followed by hydrolysis. Delignification and hydrolysis use similar compounds or enzymes. Cellulose hydrolysis enzymatically produces slightly higher bioethanol content than hydrolysis using acidic compounds (Palmqvist & Hahn-Hagerdal, 2000). An enzymatic process is highly costly. Therefore, recycling and recovering cellulase enzymes are strongly needed to reduce the high cost of production (Iranmahboob et al., 2002). Acid concentration during hydrolysis and reaction temperature are important variables. Acid compounds that are commonly used during hydrolysis are H_2_SO_4_ and HCl (Sun & Cheng, 2002; Orchidea et al, 2010). The results of Heriawati & Asdar’s research (2009) showed that cellulose hydrolysis of sago flour used 12% of HCl at an optimum condition (at 120-160°C). In addition, Irawan & Arifin (2012) found 12% 4M HCl in hydrolysis of organic waste cellulose (at 120 °C for 45 minutes).

Hydrolysis using enzymes is extremely costly. Therefore, in this study, an acid compound (HCl) was utilized. Sago waste involved in the bioethanol production was in the form of wet solid, dry solid, and liquid waste that were fermented by baker’s and tapai yeast. Research on producing bioethanol from sago starch was carried out by Nugraha (2012) who used alpha amylase and glucoamylase enzymes, and yeast. In addition, sago pulp was utilized by Gustari et al (2012) to produce bioethanol with comparative treatments of sago pulp and tapai yeast. The fundamental difference between this study and the previous research lies on the type of sago waste (wet solid waste, dry solid waste, and liquid waste) and the type of yeast (bread yeast and tapai yeast) used in bioethanol production. Parameters measured in this study which include bioethanol production levels, bioethanol profiles, and the effect of waste and yeast types on bioethanol levels have not been revealed in the previous research.

## Materials and Methods

The equipment used in the experiment included a set of glassware (pyrex), a set of fermentation equipment, a set of distillation equipment, a mortar and a pestle, a blender, an oven, a sieve (50 mesh), an autoclave, a water bath, a centrifuge, a refractometer, a pH indicator, a methylated spirit burner, an analytical balance, a filter funnel, and Gas Chromatography (GC-MS). The materials used were liquid and solid sago waste, glucose, reagent miller (DNS), aquadest, K2Cr2O7, H2SO4, urea with pro-analyst grade made by Merck, ammonium sulfate, HCl pa 37% Merck, NaOH Merck, ethanol absolute Merck, baker’s yeast, and tapai yeast. The research procedures are described as follows:

### Sample Preparation

Coarse fibers of wet solid and dry solid sago waste were cut into long, uniform strips, then put into an oven (120 °C) for 4 hours. After dried, they were cut into small pieces and blended to powder. The powder was sieved using a 50-mesh sieve. Powder that passed through the sieve was used as the sample of the research.

### Delignification of Sago Waste Powder

A hundred grams of dry-solid sago waste powder was added with 1400 ml NaOH solution 0.01 M,, heated at 90 °C, and stirred continuously. The solution was filtered using filter paper. The separated powder was rinsed with water at 100 °C (pH 7), and roasted at a temperature of 100-105 °C for four hours. Then, the powder was blended. The result of the delignifcation process was ready to be used in the next stage.

Wet solid sago waste was grinded and weighed. A hundred grams of wet solid sago waste powder was added with 1400 ml NaOH solution 0.01 M, heated at 90 °C, and stirred continuously for 60 minutes. The solution was let cool at the room temperature and then filtered. The filtered sample was then used in the hydrolysis process. Liquid sago waste was obtained from sago processing centers. The collected waste was used in hydrolysis without delignification.

### Hydrolysis

Sixty grams of (wet and dry solid) sago waste powder taken from the delignification stage was added with HCl 12% as the catalyst until 100 ml solution was obtained. The solution was poured into a three-neck flask with condenser, heated at 90 °C for 60 minutes. Sixty milliliters of liquid sago waste was hydrolyzed using 40 ml HCl 12%. The hydrolysis result was fermented with pH 5.

### Fermentation

A hundred milliliters of filtrate from the hydrolysis process was put into a container and added with 4M NaOH until its pH reached 5. It was then added with 6 grams of ammonium sulfate and 6 grams of urea as nutrients and pasteurized at 120 °C for 15 minutes then let cool. Before yeast was added, the solution was induced with 1gr/L sucrose. A hundred grams of baker’s yeast or tapai yeast was added for each treatment. Next, incubation was carried out by tightly closing the container and connecting the hose of the container to another container that contains water at a temperature of 27-30 °C for 14 days. The filtrate was used in the distillation process (Retno and Nuri, 2011).

### Distillation

Fermented filtrate was distilled with the addition of boiling stones at 80 °C to produce higher bioethanol content and then analyzed qualitatively using K_2_Cr_2_O_7_ reagent. The level and profile of bioethanol produced from each treatment were measured using a refractometer and Gas Chromatography (GC).

### Qualitative Assay of Bioethanol Using K_2_Cr_2_O_7_ as the Reactor

To identify the presence of bioethanol in fermented filtrate, a chemical qualitative test was conducted using K_2_Cr_2_O_7_. Two milliliters of K_2_Cr_2_O_7_ solution was put into 2 different test tube. Each tube was added with 5 drops of concentrated H_2_SO_4_ and shaken until homogeneous. The first tube was added with 1 ml of bioethanol to expect a change in color from orange to green or blue, whereas the second tube was added with 1ml of fermented filtrate. Filtrate containing bioethanol would change its color from orange to green or blue.

### Quantitative Assay of Bioethanol Using Refractometer

Bioethanol levels from the distillation results were analyzed using a refractometer. Bioethanol content was expressed in a refractive index value and converted into percentage. Bioethanol produced from baker’s yeast and tapai yeast fermentation was purified using a distillation device. The distillation result was accommodated in an erlenmeyer flask and dropped on the tip of the refractometer to measure its refractive index value. The refractive index value was then converted into a percentage unit based on the ethanol conversion table.

### Bioethanol Profile

GCMS (Gas Chromatography Mass Spectroscopy) analysis was performed by reading spectra on GC and MS instruments. Sample that was composed of numerous compounds will show plenty of peaks on GC spectra. Thus, based on data on retention times acquired from related literature, compounds contained in a sample can be identified. The suspected compound was then inserted into the mass spectroscopy instrument. This was done because one of the functions of gas chromatography is to separate compounds from a sample. The results were obtained from mass spectroscopy spectra on different graphs.

The qualitative data of this study were collected in the form of bioethanol profiles that were analyzed qualitatively using GCMS. GCMS analysis resulted in bioethanol graphs that were described based on peaks shown by each treatment sample. In addition, the quantitative data of this study were gathered in the form of bioethanol content. These data were analyzed qualitatively using a refractometer. Besides bioethanol content, the effect of sago waste and yeast types on bioethanol content was also described in the findings.

## Findings

### Bioethanol Profiles of Distilled Fermented Sago Waste

Glucose resulting from the hydrolysis process was fermented anaerobically by baker’s and tapai yeast for 14 days. After 14 days, the fermentation results were distilled. The distillation results were further analyzed using gas chromatography. GC chromatogram figures of the distilled fermented sago waste are presented in Figure 1.

**Figure 1.**
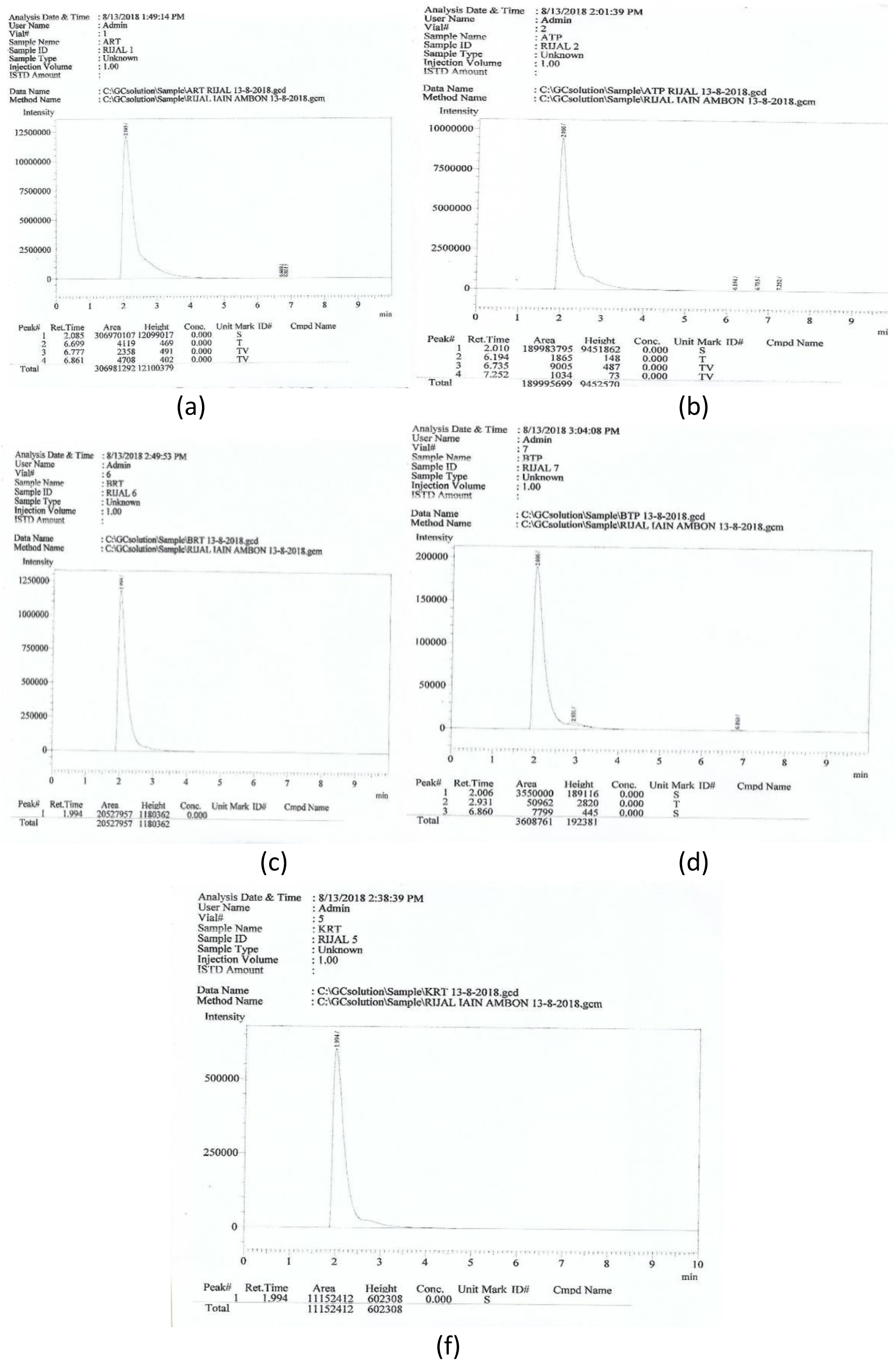
GC Chromatogram Figures of the Distilled Fermented Sago Waste (a) Liquid Sago Waste + Baker’s Yeast, (b) Liquid Sago Waste + Tapai Yeast, (c) Wet Solid Sago Waste + Baker’s Yeast, (d) Wet Solid Sago Waste + Tapai Yeast, (e) Dry Solid Sago Waste + Baker’s Yeast, and (f) Dry Solid Sago Waste + Tapai Yeast

The GC chromatogram figures were used to calculate bioethanol production levels at varied fermentation periods by comparing the sample chromatogram area with standard ethanol table. Ethanol p.a has 100% purity and a retention time of 1.944. Chromatogram of standard ethanol is shown in Figure 2.

**Figure 2.**
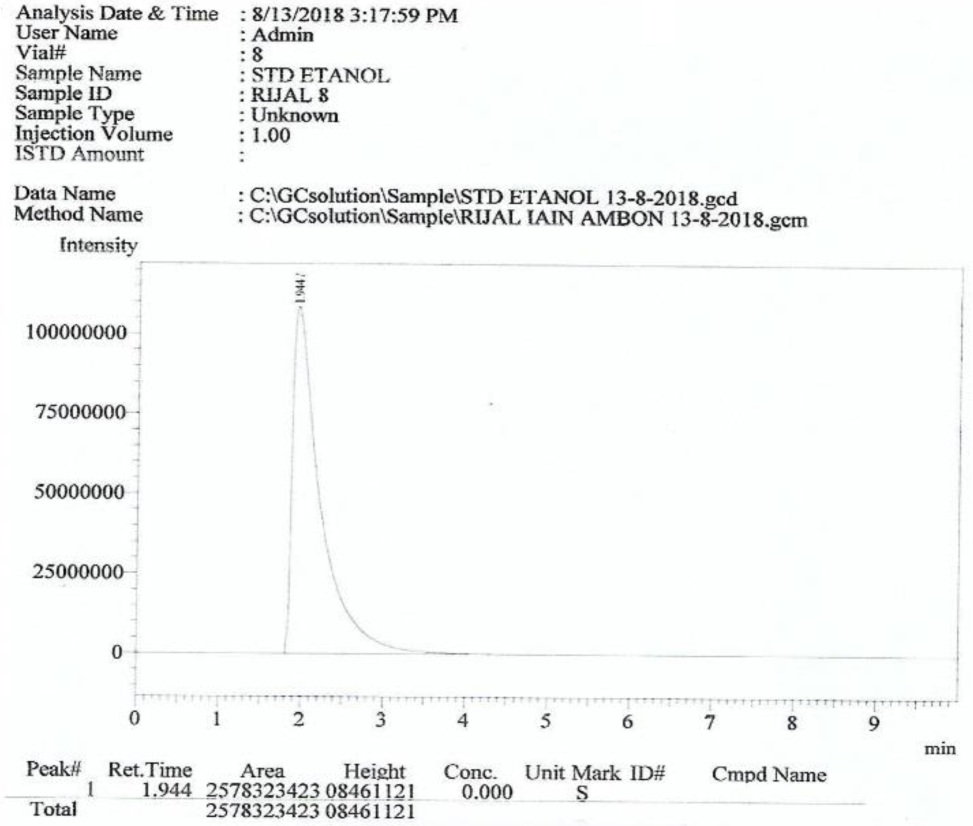
Ethanol p.a.

### Bioethanol Levels of Distilled Fermented Sago Waste

Wet solid, dry solid, and liquid sago waste contains carbohydrates in the form of lignocellulose that can be broken down into glucose through hydrolysis using acidic and basic compounds. Glucose produced during hydrolysis was fermented anaerobically by baker’s yeast and tapai yeast into bioethanol. Bioethanol level produced from each treatment can be seen in Table 1.

**Table 1.** Bioethanol Level of Each Treatment.

| Code | Refractive Index | Alcohol Content (% v/v) |
| --- | --- | --- |
| ART | 1.3391 | 14.5674 |
| ATP | 1.3379 | 12.014n2 |
| BRT | 1.3537 | 45.7021 |
| BTP | 1.3327 | 0.9504 |
| KRT | 1.3333 | 2.2270 |
| KTP | 1.3331 | 1.8014 |

Table 1 shows that the lowest level of bioethanol was found in the combination of dry solid sago waste and tapai yeast, while the highest was obtained from the combination of wet sago waste and tapai yeast. Bioethanol levels produced from each treatment were 14.5674%; 12.0142%; 45.7021%; 0.9504%; 2.2270%; and 1.8014%, respectively.

### 4.3. The Effect of Sago Waste Type and Yeast Type on Bioethanol Production Levels

The effect of sago waste type and yeast type on chromatogram area and bioethanol level of each distilled fermented waste (compared to the standard ethanol) was recorded in Table 2.

**Table 2.** The Effect of Sago Waste Type and Yeast Type on Bioethanol Levels.

| Code | Retention Time | Area | Vol (ml) | Bioethanol Content (%) |
| --- | --- | --- | --- | --- |
| ART | 2.085 | 3069.70107 | 1.00 | 14.5674 |
| ATP | 2.010 | 1899.83795 | 1.00 | 12.0142 |
| BRT | 1.994 | 205.27957 | 1.00 | 45.7021 |
| BTP | 2.006 | 355.00000 | 1.00 | 0.9504 |
| KRT | 1.994 | 111.52412 | 1.00 | 2.2270 |
| KTP | 2.095 | 361.25148 | 1.00 | 1.8014 |

The highest bioethanol level was obtained from the combination of wet solid sago waste and tapai yeast fermentation that lasted for 14 days. This condition, according to Ariyani (2012), existed because during the fermentation process, microbial activity was inhibited and heading towards the death phase. Therefore, bioethanol produced was oxidized to carboxylic acids.

## Discussion

### Bioethanol Profiles of Distilled Fermented Sago Waste Based on Types of Waste and Yeast

GC chromatogram figures of distilled fermented glucose were used to identify the level of biethanol produced from each treatment. Chromatogram area of each sample was compared to the standard ethanol that had 100% purity and a retention time of 1.944. Two or more than two peaks appeared on the ART, ATP, BTP, and KTP chromatogram figures, while only one peak appeared on the BRT and KRT chromatogram figures. The number of peaks appeared on each chromatogram figure represents the number of ethanol compounds formed during the process. Since ethanol underwent oxidation to carboxylic acids or other types of compounds due to a long fermentation period, two or more than two peaks might appear on the chromatogram figures (Ariyani, 2013).

### Bioethanol Production from Sago Waste

Sago waste contains lignocellulose. According to Hambali et al. (2007), lignocellulosic materials can be converted into bioethanol that can be used as an additive in gasoline fuel for transportation purposes. A lignocellulose compound consists of three main components, namely cellulose, hemicellulose, and lignin that make up the plant cell walls (Hermiati et al., 2010). In this study, the process of converting sago waste as a lignocellulosic material into biethanol underwent three stages, namely delignification, hydrolysis, and fermentation. Hydrolysis was performed to convert cellulose into simple sugars and yeast fermentation was carried out to break down the sugars into bioethanol (Hermiati, 2010). Different types of sago waste contain different levels of lignocellulose. Therefore, there was an effect of types of sago waste used in this study on bioethanol produced. The highest level of bioethanol found in this study (45.7021%) was found in the combination of wet solid waste and baker’s yeast fermentation. This figure is higher than those found by Nugraha (2012) (1.110%) and Gustari et al (2012) (2.1%).

### The Effect of Sago Waste and Yeast Types on Bioethanol Content

Sago waste, either in a solid or a liquid form, is a carbon source that can be used by microorganisms to grow in a growth medium (Hambali et al., 2007). Solid and liquid waste has different chemical structures and carbohydrate content. Solid waste contains higher carbohydrates than liquid waste because carbohydrates in liquid waste can be easily dissolved into the environment. The combination of types of waste and types of yeast resulted in different levels of bioethanol. The treatment that produced the highest bioethanol content (45.7021%) was the combination of wet solid waste + baker’s yeast (BRT), while the treatment that produced the lowest bioethanol content (0.9504%) was the combination of wet solid waste + tapai yeast (BTP). Wet solid sago waste contains high carbohydrate content. Carbohydrates found in such type of waste will result in high amount of glucose after hydrolysis (Hermiati et al., 2010). Glucose will be fermented by yeast into alcohol under an anaerobic condition (Ari and Hadi, 2008; Azizah N, 2012; Taherzadeh & Karimi, 2007). Baker’s yeast contains a bacterium, known as *S. Cereviceae.* This bacterium can support the optimization of glucose-ethanol fermentation (Retno & Nuri, 2011; Santi, 2008; Afriani, 2011). Optimal fermentation processes will result in higher alcohol content (Aziza, 2010).

## Conclusion

1. Solid and liquid sago waste is lignocellulosic biomass that is composed of carbohydrate components. Lignocellulose can be converted into glucose through hydrolysis using an acid compound (such as 12% HCL). Glucose fermented by baker’s yeast for 14 days at a temperature of 27 – 30 ° C under an anaerobic condition produced bioethanol with different levels. The combination of wet solid sago waste and baker’s yeast fermentation produced the highest level of bioethanol (45.70%) compared to other treatments.
2. Gas chromatography in this study was used to identify compounds contained in a sample. The number of peaks shown on a chromatogram figure indicates the number of compounds contained in a sample. Based on the gas chromatography analysis, it was found that the ART, ATP, BTP, or KTP treatment contained two or more than two compounds, while the BRT or KRT treatment contained one compound, indicated by the appearance of 1 peak on their chromatogram figures.
3. The results of this study showed that types of sago waste and types of yeast had an effect on bioethanol production levels. The highest level of bioethanol was obtained from the combination of wet solid sago waste and baker’s yeast fermentation. Wet solid sago waste contains lignocellulose, while baker’s yeast contains *S. cereviceae*, a microorganism that has the ability to convert glucose into ethanol under an anaerobic condition.

